# Heliorhodopsins are absent in diderm (Gram-negative) bacteria: Some thoughts and possible implications for activity

**DOI:** 10.1101/356287

**Authors:** José Flores-Uribe, Gur Hevroni, Rohit Ghai, Alina Pushkarev, Keiichi Inoue, Hideki Kandori, Oded Béjà

## Abstract

Microbial heliorhodopsins are a new type of rhodopsins with an opposite membrane topology compared to type-1 and −2 rhodopsins, currently believed to engage in light sensing. We determined their presence/absence is monoderms and diderms representatives from the *Tara* Oceans and freshwater 25 lakes metagenomes. Heliorhodopsins were absent in diderms, confirming our previous observations in cultured Proteobacteria. Based on these observations, we speculate on the putative role of heliorhodopsins in light-driven transport of amphiphilic molecules.

---

Many organisms capture or sense sunlight using rhodopsin pigments (Spudich et al., 2000; Ernst et al., 2014). Rhodopsins are currently divided to two distinct protein families (Spudich et al., 2000): type-1 (microbial rhodopsins) and type-2 (animal rhodopsins). The two families share similar topologies with 7 trans-membrane (TM) helices, however, the two families show little or no sequence similarity to each other. Recently, using functional metagenomics, another family of rhodopsins, the heliorhodopsins, was detected (Pushkarev et al., 2018). The orientation of heliorhodopsins in the membrane is opposite to that of type-1 or type-2 rhodopsins, with the N-terminus facing the cell cytoplasm. Heliorhodopsins show photocycles longer than 1 second, suggestive of light sensory activity.

Heliorhodopsins are found in diverse bacteria, archaea, unicellular eukarya and viruses (Pushkarev et al., 2018). These microbes range from psychrophiles, mesophiles and even hyperthermophiles originating from soil, freshwater, marine and hypersaline environments. Most peculiarly, the new family is completely missing in cultured proteobacteria (Pushkarev et al., 2018), a diderm phylum in which numerous type-1 rhodopsins are detected (DeLong and Béjà, 2011; Pinhassi et al., 2016). In order to confirm the absence of heliorhodopsins in diderms, we have searched for heliorhodopsins in *Tara* Oceans (Brum et al., 2015; Sunagawa et al., 2015), several freshwater lakes metagenomes, the UBA collection of 7,903 metagenome assembled genomes (MAGs) from multiple habitats (Parks et al., 2017), as well as in the Universal Protein Resource (UniProt) database.

We scanned a non-redundant collection of all the open reading frames from the *Tara* Oceans microbiome and virome as well as metagenomic freshwater lakes datasets to identify putative rhodopsins using hidden Markov Models (HMM) through graftM (Boyd et al., 2018; Potter et al., 2018). Putative rhodopsins where then analyzed using a HHsuite pipeline based on HHPred for structure prediction homology search (Söding, 2005; Zimmermann et al., 2017) against the Protein Data Bank (PDB) structures database (see methods). Since the HHsearch probability is recommended as the principle measure of statistical significance (Söding, 2005; Zimmermann et al., 2017), the conventional probability threshold for determining homology is usually set above 80-90%. Interestingly, the probability of most of the rhodopsin containing contigs (i.e. type-1 and heliorhodopsin) is above the conventional threshold, without the ability to differentiate rhodopsins by type (Fig. 1a, right pane). This can explain why some of the heliorhodopsins found in public databases are annotated as “rhodopsin-like” proteins. In contrast, the HHsearch score does show two distinct groups: type-1 and heliorhodopsin (Fig. 1a, top pane). From the distribution of probability vs. score (Fig. 1a center pane), it seems that beyond rhodopsins assigned to type-1 or heliorhodopsins no other rhodopsin groups are clearly observed. This might suggest that microbial rhodopsins are limited to these two groups only. Furthermore, no heliorhodopsin were affiliated with proteobacteria or to any other diderms in metagenomes (Fig. 1b) or UniProt (Fig. S1), confirming our previous observation with cultured proteobacteria representatives (Pushkarev et al., 2018). Interestingly, we identified more heliorhodopsins than type-1 rhodopsins in viruses. The only exception to the restriction of heliorhodopsins to monoderms was the genome of a Sphaerochaeta MAG UBA6084 assembled from a waste-water metagenome. No other Sphaerochaeta MAGs or complete genomes encoded any heliorhodopsins. While Sphaerochaeta are diderms, they appear to have a history of massive horizontal gene transfers, nearly 40% of genes in their genomes appear to originate from clostridia (monoderms) (Caro-Quintero et al., 2012). They also have an atypical cell wall structure (reduced cross-linking in peptidoglycan owing to absence of penicillin-binding protein PBP), coupled with the lack of characteristic axial flagella that are found in all Spirochaetes leading to a non-rigid cell walls and spherical morphologies in contrast to the other members of the phyla.

**Figure 1.**
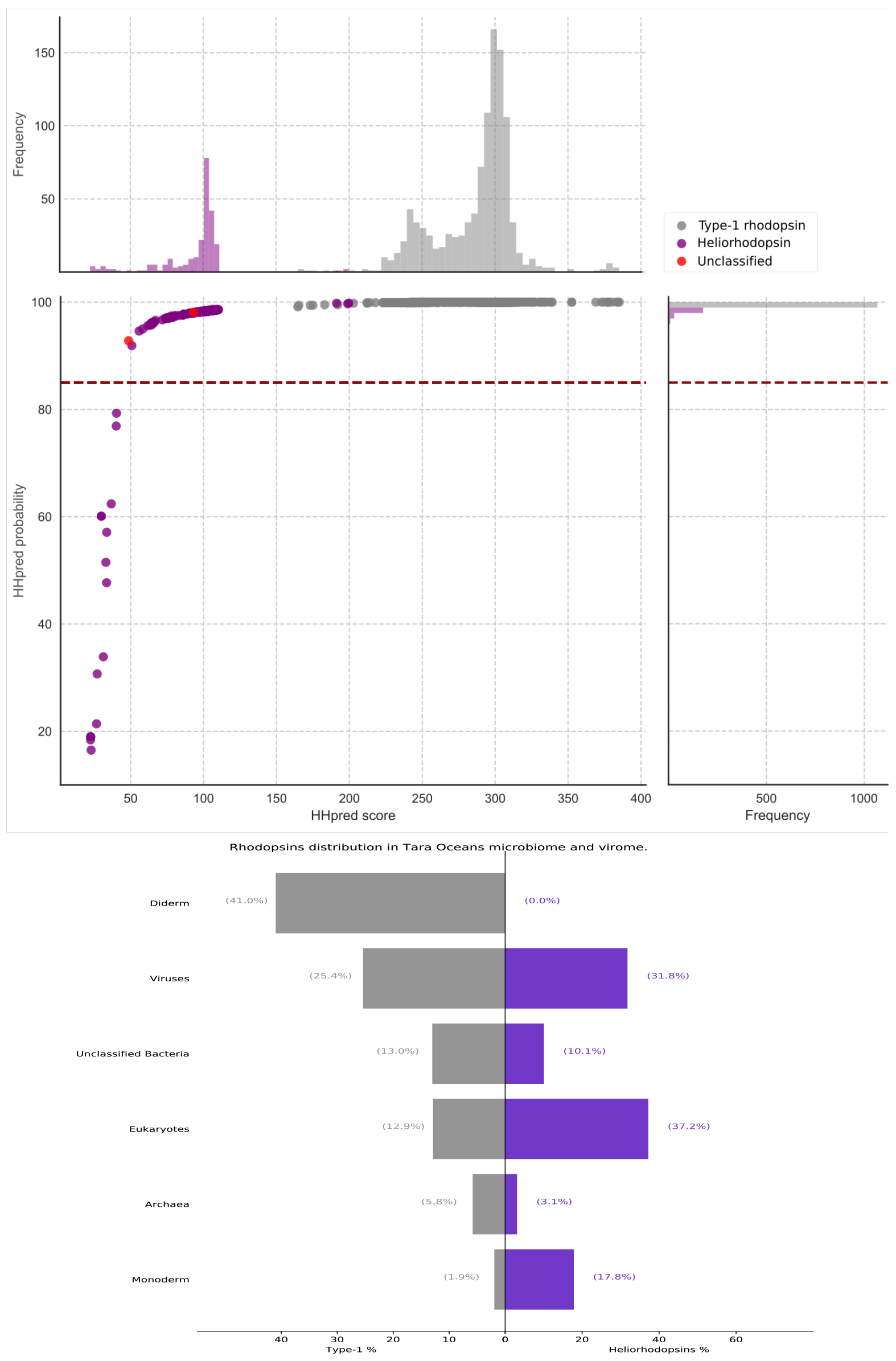
Heliorhodopsins and type-1 rhodopsins in *Tara* Oceans’ diderms and monoderms. (**a**) Structural-based prediction of putative rhodopsins using HHsearch probability and score values. Absence of heliorhodopsins structures in the PDB reflect in the low HHsearch score and probability compared to the type-1 rhodopsins. (**b**) Presence/absence of heliorhodopsins in diderms (Proteobacteria, Aquificae, Chlamydiae, Bacteroidetes, Chlorobi, Cyanobacteria, Fibrobacteres, Verrucomicrobia, Planctomycetes, Spirochetes, Acidobacteria), monoderm (Actinobacteria, Firmicutes, Thermotogae, Chloroflexi), Archaea, Eukarya and viruses. Separation to diderms and monoderms is according to Gupta 2011 (Gupta, 2011).

We suggest several possible scenarios to explain the absence of heliorhodopsins in diderms: (*i*) Heliorhodopsins evolved in monoderm (Gram-positive) bacteria only. As lateral gene transfer was shown to be ubiquitous for type-1 rhodopsins (Frigaard et al., 2006) and assuming similar evolutionary forces are at work on heliorhodopsins, this explanation seems less likely. (*ii*) Absence of the outer membrane in monoderm bacteria. The outer membrane of diderm bacteria is a semi-permeable barrier and can block passage of different amphiphilic compounds (Egan, 2018) or light if heavily pigmented (Kumagai et al., 2018). This might suggest the possible involvement of heliorhodopsins in light dependent transport of such compounds in monoderms (Fig. 2).

**Figure 2.**
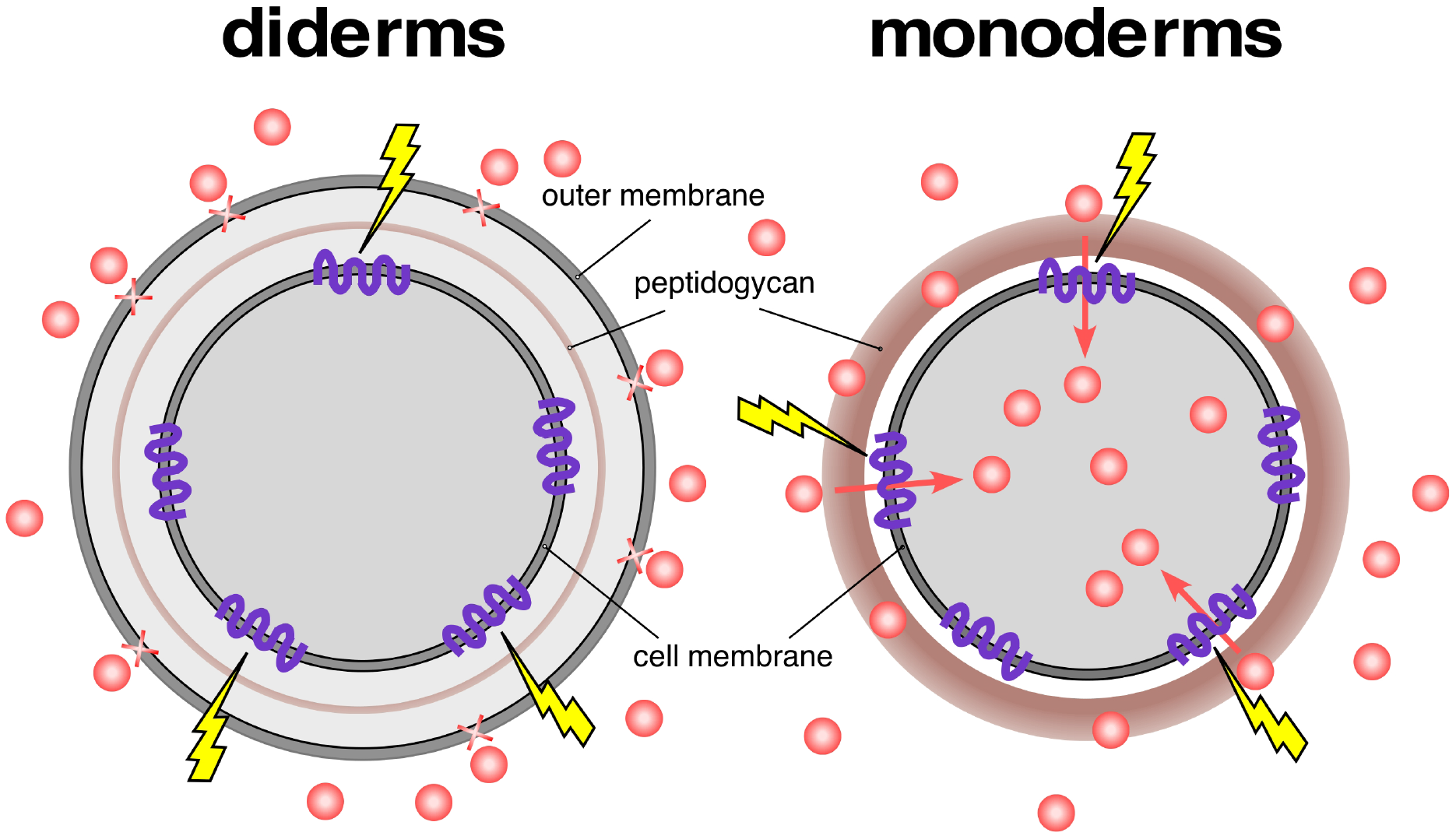
Suggested role for heliorhodopsins in light-driven transport of amphipilic compounds. Amphiphilic molecules do not pass the outer membrane in diderms and therefore are not available to transport by heliorhodopsins. Amphiphilic molecules are depicted in red and heliorhodopsins in purple.

## Concluding remarks

The function of heliorhodopsins is currently unknown. Based on their very slow photocycle (> 1 sec) we recently suggested that heliorhodopsins act as sensory rhodopsins. Their complete absence in diderms, a group containing numerous type-1 rhodopsin families, potentially points to heliorhodopsins involvement in light-driven transport of molecules that cannot pass the outer membrane of diderms (*e.g.* amphiphilic molecules). Future experiment with cultured monoderms containing heliorhodopsins or with permeable outer membrane *Escherichia coli* mutants (Béjà and Bibi, 1996) might help in resolving this issue.

## Acknowledgements

The authors thank Sheila Roitman for the initial observation and inspiration. This work was supported by Israel Science Foundation - F.I.R.S.T. (Bikura) Individual grant (545/17).

